# Detecting M/EEG modular brain states in rest and task

**DOI:** 10.1101/510727

**Authors:** A. Kabbara, M. Khalil, G. O’Neill, K. Dujardin, Y. El Traboulsi, F. Wendling, M. Hassan

## Abstract

The human brain is a dynamic networked system that continually reconfigures its connectivity patterns over time. Thus, developing approaches able to adequately detect fast brain dynamics is critical. Of particular interest are the methods that analyze the modular structure of brain networks, i.e. the presence of clusters of regions that are densely inter-connected. In this paper, we propose a novel framework to identify fast modular states that dynamically fluctuate over time during rest and task. We validated our method using MEG data recorded during a finger movement task, identifying modular states linking somatosensory and primary motor regions. The algorithm was also validated on dense-EEG data recorded during picture naming task, revealing the sub-second transition between several modular states which relate to visual processing, semantic processing and language. Next, we validated our method on a dataset of resting state dense-EEG signals recorded from 124 patients with Parkinson’s disease and different cognitive phenotypes. Results disclosed brain modular states that differentiate cognitively intact patients, patients with moderate cognitive deficits and patients with severe cognitive deficits. Our new approach tracks the brain modular states, in healthy subjects and patients, on an adequate task-specific timescale.

## 1. Introduction

The human brain is a modular dynamic system. Following fast neuronal activity (Pfurtscheller and Aranibar, 1977; Pfurtscheller and Lopes Da Silva, 1999), the functional organization of resting (Baker et al., 2014; Damaraju et al., 2014; de Pasquale et al., 2015, 2012; Kabbara et al., 2017) and task-evoked connectivity (Bola and Sabel, 2015; Hassan et al., 2015; Hutchison et al., 2013; O’Neill et al., 2017b) are in constant flux. Hence, an appropriate description of time-varying connectivity is of utmost importance to understand how cognitive and behavioral functions are supported by networks.

Electro/magneto-encephalography (EEG/MEG) are unique noninvasive techniques, which allow for the tracking of brain dynamics on a millisecond time-scale, a resolution not reachable using other techniques such as the functional Magnetic Resonance Imaging (fMRI) (Cohen, 1972; Nunez and Srinivasan, 2007; Penfield and Jasper, 1954). In this context, several methods have been proposed to reveal when, and how the functional connections between brain regions vary during short-time (sub-second) experiments. Some of these studies proposed to group the temporal networks into states, where each state reflect unique spatial connectivity pattern. These brain states were mainly generated using Hidden Markov model approaches (Baker et al., 2014; Vidaurre et al., 2017), K-means clustering (Allen et al., 2017, 2014; Damaraju et al., 2014) or independent component analysis (O’Neill et al., 2017b). Other studies have tried to investigate the dynamic topological changes using graph theoretical analysis (de Pasquale et al., 2015; Kabbara et al., 2017).

Due to the modular organization of the human brain network (Sporns and Betzel, 2016a), methods for detecting network communities (or modules) are of particular interest (Sporns and Betzel, 2016b). These methods decompose the network into building blocks or modules that are internally strongly connected, often corresponding to specialized functions. Importantly, brain modularity has been revealed to be related to behavior involved in learning (Bassett et al., 2011), remembering, attention and integrated reasoning (Gallen et al., 2016).

To detect modules, many community detection methods have been proposed (M. Girvan and Newman, 2002; Newman and Girvan, 2004). Using the most widely applied community detection methods, brain networks are often analyzed as static arrangements of nodes and edges with a number of studies starting to explore how the modular organization shapes the dynamics of neural activity (Bassett et al., 2015a, 2013, 2011; Sporns and Betzel, 2016b). One of the algorithms used to characterize the flow of communities across time is multi-slice modularity (also called multi-layer modularity) (Mucha et al., 2010), which has been used to follow changes of the modular architecture across time (Bassett et al., 2015b, 2011). However, using this technique solely, there are basically two strategies to analyze the modular structures: i) generate a community structure consistent over time which will constrain the dynamic analysis or ii) consider each temporal community structure as a state which is technically very difficult and doesn’t reflect the underlying brain functions. Hence, it does not ‘automatically’ decipher functional modular brain states i.e. subset of brain modules implicated in a given brain function at a given time period.

Here, we propose a novel framework aiming to elucidate the main modular brain structures, called ‘modular states’ (MS), which fluctuate over time during rest and task. This new method is based on temporally categorizing the modular structures that share the same topology by quantifying the similarity between their partitions (Figure 1). The proposed framework was validated in simulation and on three different MEG/EEG datasets, recorded in rest and task from healthy and/or patients. Our method revealed both time-varying functional connectivity and the corresponding spatial patterns. We were able to track the space/time dynamic of brain networks in task-adapted timescale.

**Figure 1.**
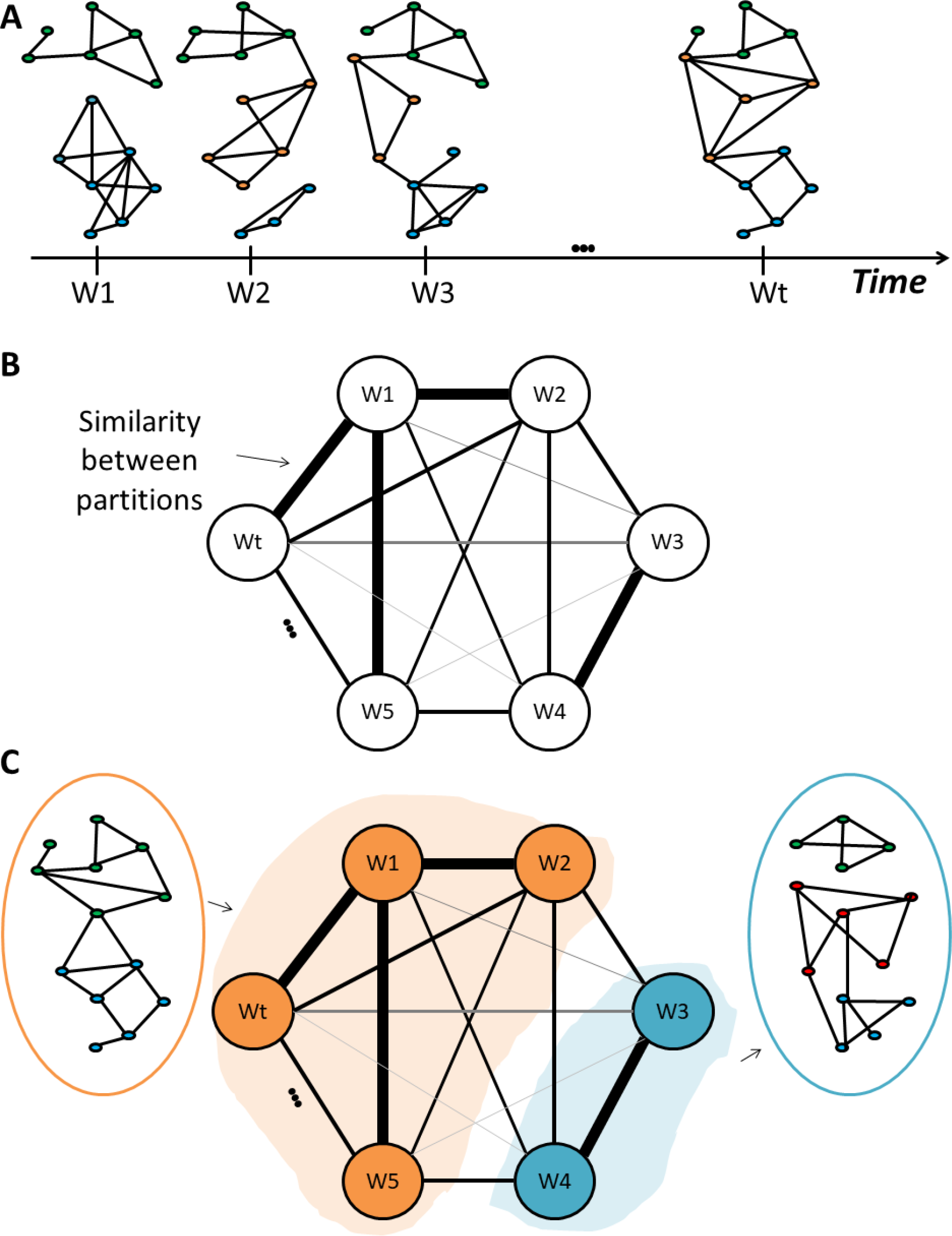
The algorithm procedure. A) Computation of modules for each temporal network (Wt). B) Assessment of the similarity between the dynamic modular structures. C) Clustering the similarity matrix into “categorical Modules”.

## 2. Materials and methods

We have developed two versions of the algorithm i) “categorical” where we aim to find the main modular structures over time, with no interest in their sequential order and ii) “consecutive” where the objective is to find the modular structures in a successive way. The two versions are described hereafter.

### 2.1. Categorical version

It includes four main steps:

1. Compute the dynamic functional connectivity matrices between the regional time-series using a sliding window approach (Allen et al., 2014; Hutchison et al., 2013; Kabbara et al., 2017). Consequently, a weighted network is generated at each time window (Figure 1. A).
2. Decompose each network into modules (i.e clusters of nodes that are internally strongly connected, but externally weakly connected) (Figure 1. A). To do that, different modularity algorithms were proposed in the literature (Blondel et al., 2008b; Duch and Arenas, 2005; M. Girvan and Newman, 2002; Guimerà and Amaral, 2005). In our study, we adopted the consensus clustering approach which was previously used in many studies (Bassett et al., 2013; Kabbara et al., 2017): Given an ensemble of partitions acquired from the Newman algorithm (M Girvan and Newman, 2002) and Louvain algorithm (Blondel et al., 2008a) repeated for 200 runs, an association matrix is obtained. This results in a N*N matrix (N is the number of nodes) and an element *A*_*i,j*_ represents the number of times the nodes *i* and *j* are assigned to the same module across all runs and algorithms. The association matrix is then compared to a null model association matrix generated from a permutation of the original partitions, and only the significant values are retained (Bassett et al., 2013). To ultimately obtain consensus communities, we re-clustered the association matrix using Louvain algorithm.
3. Assess the similarity between the temporal modular structures (Figure 1. B). In this context, several methods have been suggested to compare community structures (Traud et al., 2008). Here, we focused on the pair-counting method, which defines a similarity score by counting each pair of nodes drawn from the N nodes of a network according to whether the pair falls in the same or in different groups in each partition (Traud et al., 2008). We considered the z-score of Rand coefficient, bounded between 0 (no similar pair placements) and 1 (identical partitions). This yield a T*T similarity matrix where T is the number of time windows.
4. Cluster the similarity matrix into “categorical” modular states (MS) using the consensus clustering method (Figure 1. C). This step combines similar temporal modular structures in the same community. Hence, the association matrix of each “categorical” community is computed using the modular affiliations of its corresponding networks.

### 2.2. Consecutive version

The difference between the two versions of the algorithm is essentially in the fourth step, in which the final communities were defined. Particularly, the similarity matrix is segmented in a sequential way using the following steps:

- Threshold the similarity matrix using an automatic thresholding algorithm described in (Genovese et al., 2002). Briefly the matrix was converted into a *p*-value map which is then thresholded based on the false discovery rate (FDR) controlling.
- Apply a median filter in order to get smoother presentation of the similarity matrix.
- Segment the matrix in a sequential way following the algorithm illustrated in the flowchart of Figure 2. In brief, the method groups similar consecutive modular structures. As these modular structures show high similarity values with each other, the algorithm detects the squares located around the diagonal of the similarity matrix. As presented in Figure 2, the condition for which two consecutive structures are associated to the same state is the following:

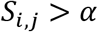

Where 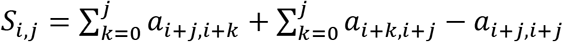; *i*, *j ϵ* [1, *T*] where *a*_*l,m*_ denotes the similarity value between the modular structure corresponding to the time window *l* and that corresponding to the time window *m*. *α* is the “accuracy parameter”, strictly bounded between 0 and 1. It regulates the temporal-spatial accuracy of detected modular states. We recommend choosing an adaptive value of *α*. In this paper, we choose *α* equals to the average of the similarity matrix. A segment is considered as relevant if the number of included time-windows is greater than *j*_*min*_ (the minimal size allowed for a segment).
- For each detected segment, the modular structure is obtained after computing the association matrix of the corresponding time windows modular affiliations.

**Figure 2.**
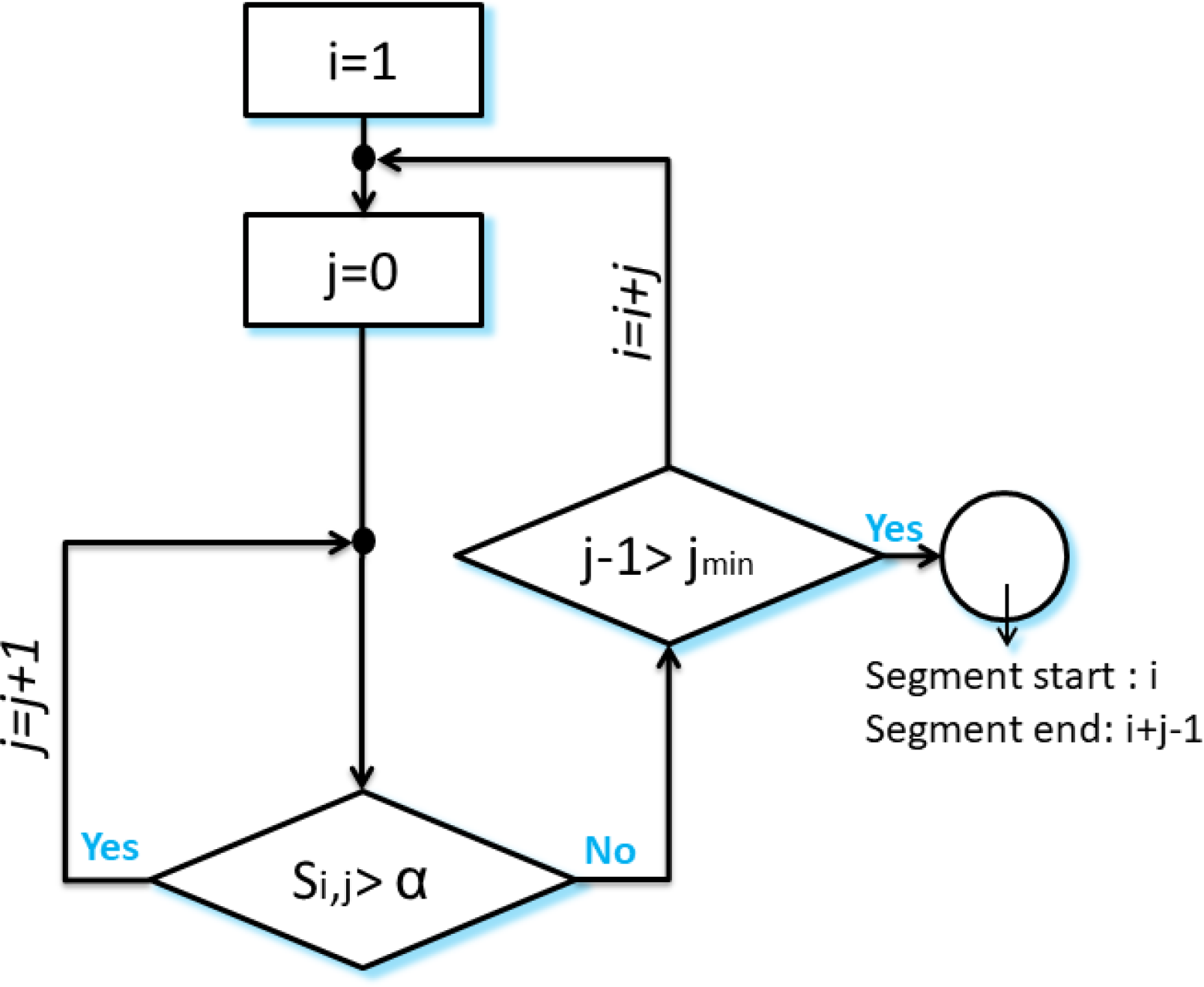
The flowchart of the segmentation algorithm. The algorithm is illustrated in Figure 3. (I): starting with *i* = 1, *j* = 0; and considering that *a*_11_ is lower than *α*, we obtained *S*_1,0_ = *a*_11_ < *α*, and the algorithm moves to the next time window *i* = 2. (II): as *S*_2,0_ = *a*_22_ is greater than *α*, *j* is incremented by 1. (III): having 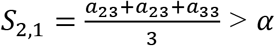, the second and the third time windows are associated to the segment. (IV): the algorithm succeeds to add also the fourth time window as 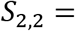 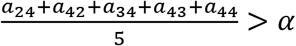. (V): Then, for *j* = 3, we obtain 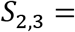 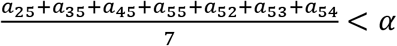. This means that the fifth time window differs from the previous windows in his modular structure. Afterwards, the algorithm moves toward finding another segment by incrementing *i* and repeating the process (IV).

**Figure 3.**
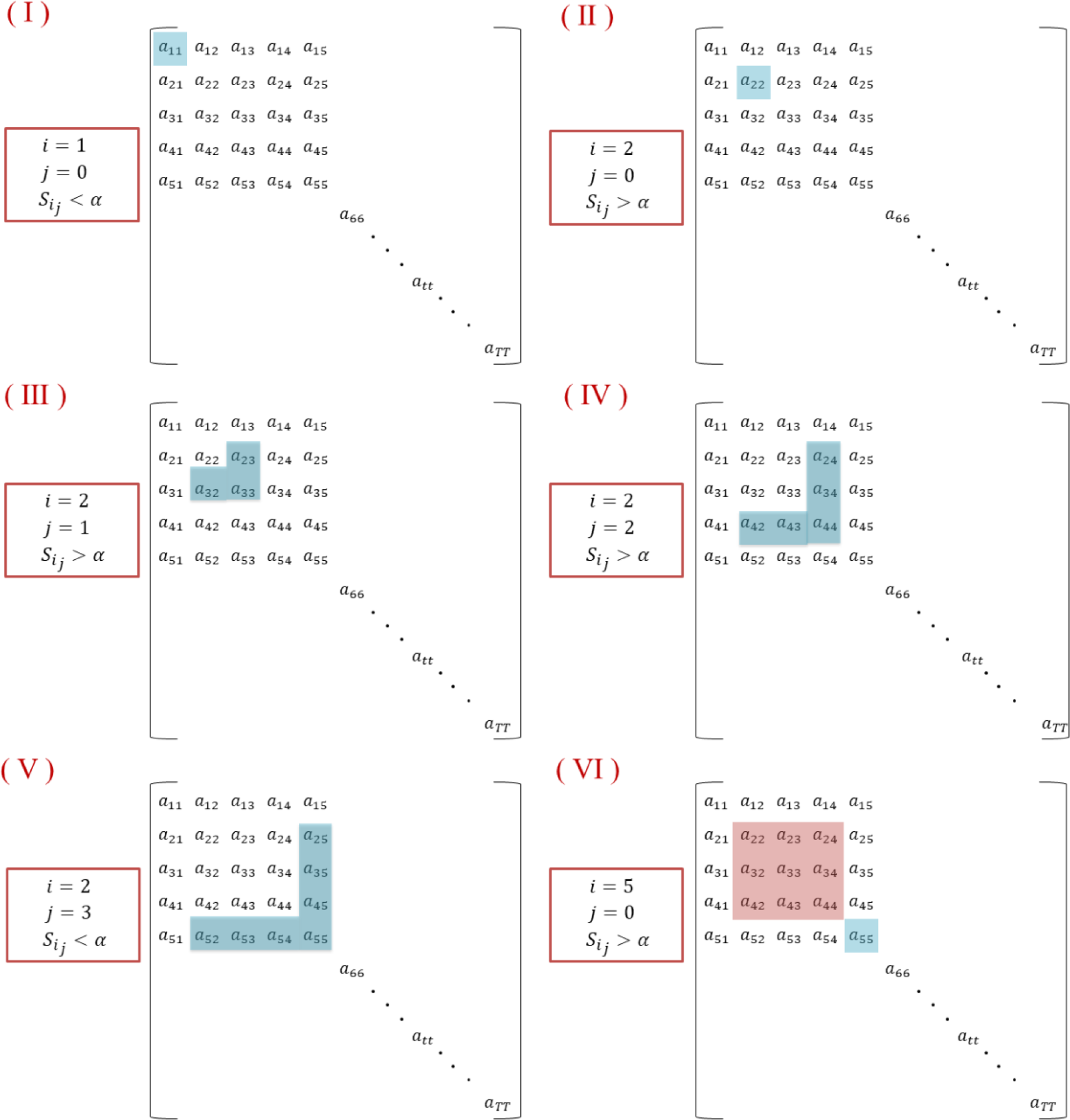
An illustrative example describing the segmentation algorithm.

### 2.3. Simulated data

We simulated adjacency tensor data following the methodology applied in (O’Neill et al., 2017b). Briefly, four N*N adjacency matrices *P*_*j*_ were constructed, where *j* ∈ [1,4] and N is the number of ROIs. We used an anatomical atlas of 221 ROIs with the mean of Desikan-Killiany atlas sub-divided by (Hagmann et al., 2008), yielding to N=221. The adjacency matrices and the 3D visualization of the networks are presented in Figure 4.A. Following this step, the time evolution of dynamic connectivity in each network is given by:

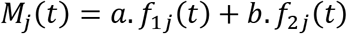

*f*_1*j*_(*t*) is the modulation function, which was represented by Hanning window of unit amplitude. *f*_2*j*_(*t*) represents uncorrelated Gaussian noise added to the simulated time-courses, and *a* and *b* are scalar values set to 0.45 and 0.15 as in (O’Neill et al., 2017b).

In our study, *M*_*j*_(*t*) is sampled at 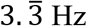 (to obtain a sliding window of 0.3s as in real data). The onset as well as the duration of each module structure is illustrated in Figure 4.B. We then combined the four network matrices in order to generate single adjacency matrix at each time point *t* over a time-course spanning 60 seconds. As a final step, we added a random Gaussian noise to the adjacency tensor, and the standard deviation of the noise was allowed to vary between 0.2 and 0.5.

**Figure 4.**
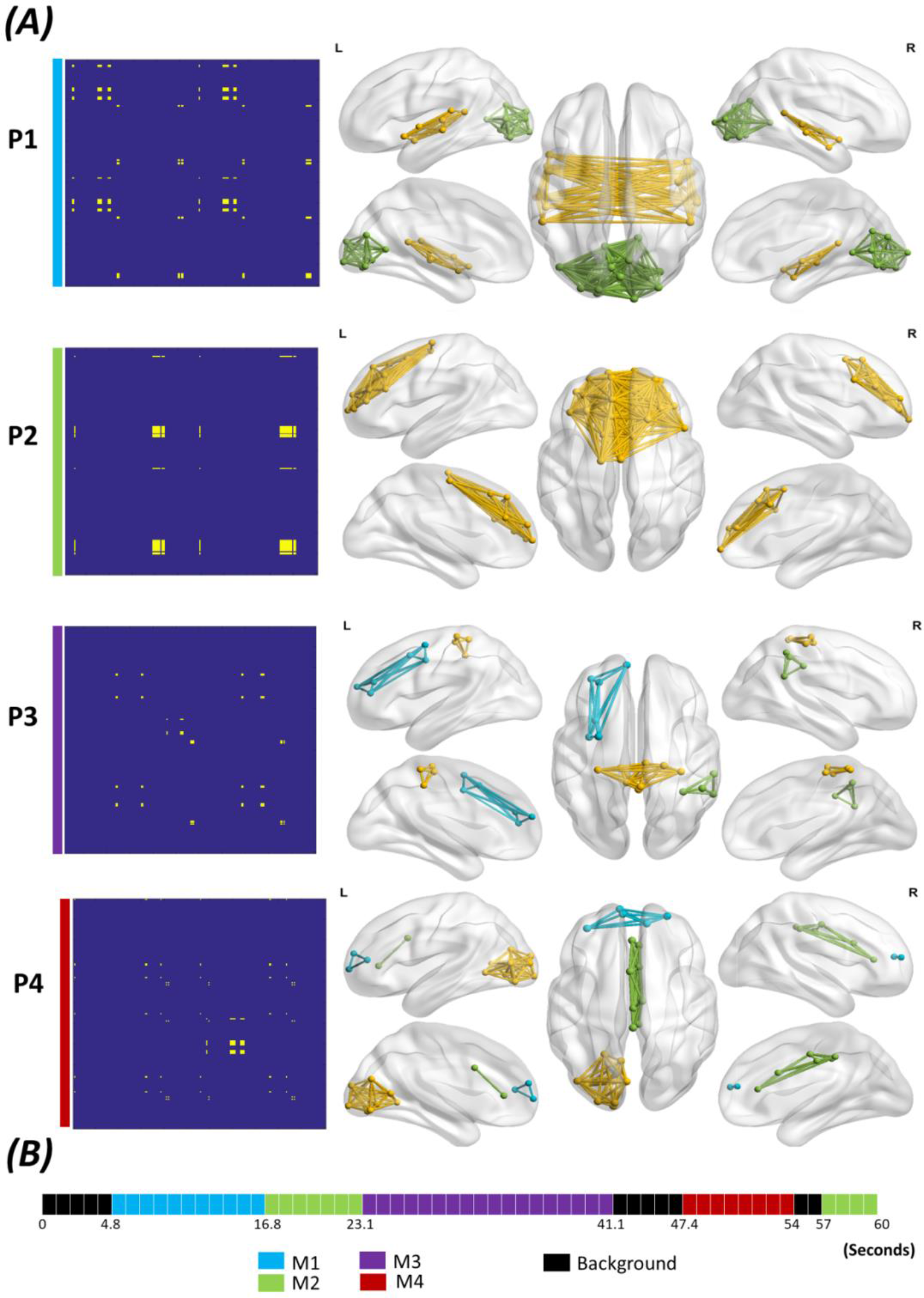
The simulation scenario. A) Left: the adjacency matrix P_j_ of the constructed networks, Right: the 3D cortical presentation of the modular structures of the simulated networks. B) the time axis showing the beginning and the end of each network M_j_ (which was generated by combining P_j_ to uncorrelated noise).

### 2.4. Validation

On simulated data, we evaluated the performance of the algorithm by computing the similarity between the reconstructed and the simulated (reference) networks, taking into account both spatial and temporal similarities. The spatial similarity is given by the z-score of Rand coefficient between the simulated and the constructed modular structures, while the temporal similarity represents the rate of the correct affiliation of time windows.

### 2.5. Real data

#### 2.5.1. Dataset 1- *Self paced motor task for healthy participants (MEG data)*

Previously used in (O’Neill et al., 2017b, 2015; Vidaurre et al., 2016), this dataset includes 15 participants (9 male, 6 female) asked to press a button using the index finger of their dominant hand, once every 30 seconds. Using a 275-channel CTF MEG system (MISL; Coquitlam, BC, Canada), MEG data were recorded at a sampling rate of 600Hz. MEG data were co-registered with a template MRI. The cortex was parcellated using the Desikan-Killiany atlas (68 regions). The pre-processing, the source reconstruction and the dynamic functional connectivity computations were performed similarly as in O’Neill et al. (O’Neill et al., 2017b). Briefly, the pre-processing comprises the exclusion of trials (t= −12s → 12s) contaminated by noise. Then, source time courses were reconstructed using a beamforming approach (please refer to (O’Neill et al., 2017b) for more details). Afterwards, the regional time-series were symmetrically orthogonalized following the method proposed in (Colclough et al., 2015) to remove the effects of “signal leakage”. The amplitude envelopes of the time courses were obtained using Hilbert transform. Finally, the dynamic connectivity was estimated by the Pearson correlation measure using a sliding window approach of 6 s of length. The sliding window was shifted by 0.5 s over time. The number of connectivity matrices obtained for each trial was then 49. The consecutive scheme of the proposed method was tested.

#### 2.5.2. Dataset 2- *Picture naming task for healthy participants (dense-EEG data)*

Twenty one right-handed healthy subjects (11 women and 10 men), with no neurological disease participated in this study. In a session of about eight minutes, each participant was asked to name 148 displayed pictures on a screen using EPrime 2.0 software (Psychology Software Tools, Pittsburgh, PA) (Schneider et al., 2002). Oral responses were recorded to set the voice onset time. This study was approved by the National Ethics Committee for the Protection of Persons (CPP), conneXion study, agreement number (2012- A01227-36), and promoter: Rennes University Hospital. All participants provide their written informed consent to participate in this study. A typical trial started with the appearance of an image during 3 sec followed by a jittered inter-stimulus interval of 2 or 3 sec randomly. Errors in naming were discarded from the analysis. A total of 2926 on 3108 events were considered.

Dense-EEG data were recorded using a system of 256 electrodes (EGI, Electrical Geodesic Inc.). EEGs were collected at 1 kHz sampling frequency and band-pass filtered between 3 and 45 Hz The pre-processing and the computation of the functional connectivity followed the same pipeline applied in (Hassan et al., 2015). In brief, each trial (t=0 → 600ms) was visually inspected, and epochs contaminated by eye blinks, muscle movements or other noise sources were rejected. As described in the previous study (Hassan et al., 2015), the source connectivity method was performed using the wMNE/PLV combination, and the dynamic functional connectivity was computed at each millisecond. Authors also used the Destrieux atlas sub-divided into 959 regions (Hassan et al., 2015). Finally, a tensor of dimension 959 − 959 − 600 was obtained and analyzed using the consecutive scheme of our algorithm.

#### 2.5.3. Dataset 3- *resting state in Parkinson′s disease patients (dense-EEG data)*

This dataset includes 124 patients with idiopathic Parkinson’s disease defined according to the UK Brain Bank criteria for idiopathic Parkinson’s disease (Gibb and Lees, 1988). These patients were separated into three groups: G1) cognitively intact patients (N=63), G2) patients with mild cognitive deficits (N=46) and G3) patients with severe cognitive impairment (N=15). All participants gave their informed consent to participation in the study, which had been approved by the local institutional review boards (CPP Nord-Ouest IV, 2012-A 01317-36, ClinicalTrials.gov Identifier: NCT01792843). Dense-EEG were recorded with a cap (Waveguard®, ANT software BV, Enschede, the Netherlands) with 122 scalp electrodes distributed according to the international system 10-05 (Oostenveld and Praamstra, 2001). Electrodes impedance was kept below 10 kΩ. Patients were asked to relax without performing any task. Signals were sampled at 512 Hz and band-pass filtered between 0.1 and 45 Hz.

The data were pre-processed according to (Hassan et al., 2017) dealing with the same dataset. Briefly, EOG artifact detection and correction was applied following the method developed in (Gratton et al) (Gratton et al., 1983). Afterwards, epochs with voltage fluctuation >+90 μV and <−90 μV were removed. For each participant, two artifact-free epochs of 40s lengths were selected. This epoch length was used previously and considered as a good compromise between the needed temporal resolution and the reproducibility of the results in resting state (Kabbara et al., 2017). To compute the dynamic functional connectivity, the steps adopted here are the same used in many previous studies (Hassan et al., 2017; Kabbara et al., 2018, 2017). First, EEG data were co-registered with a template MRI through identification of the same anatomical landmarks (left and right pre-auricular points and nasion). Second, the lead field matrix was computed for a cortical mesh with 15,000 vertices using OpenMEEG package (Gramfort et al., 2010) available in Brainstorm. The noise covariance was estimated using one minute resting segment. After that, the time-series of EEG sources were estimated using the wMNE algorithm where the regularization parameter was set according to the signal to noise ratio (λ= 0.1 in our analysis). An atlas-based segmentation approach was used to project EEGs onto an anatomical framework consisting of 68 cortical regions identified by means of Desikan-Killiany (Desikan et al., 2006) atlas. The dynamic functional connectivity was then computed using a sliding window over which PLV was calculated (Lachaux et al., 1999). In the previous study (Hassan et al., 2017), the disruptions of the functional connectivity were found in the alpha2 band (10-13 Hz). For this reason, we considered the same frequency band in our analysis. To obtain a sufficient number of cycles at the given frequency band, we chose the smallest window length that is equal to 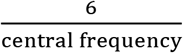 as recommended in (Lachaux et al., 1999). This yields to a sliding window of 0.52 s. We then adopted a proportional threshold of 10% to remove spurious connections from the connectivity matrices. These steps produce, for each epoch, a connectivity tensor of dimension N x N x T where N is the number of ROIs (68 regions), and T is the number of time windows (77 time-windows). This tensor is formally equivalent to dynamic functional connectivity matrices, and was analyzed using the categorical version of the proposed algorithm.

## 3. Results

The dynamic functional connectivity matrices were obtained using a sliding window approach giving a weighted network at each time window. By applying a community detection algorithm (Louvain method), each network was then decomposed into modules (i.e clusters of nodes that are internally strongly connected, but externally weakly connected). The similarity between the temporal modular structures was calculated (Figure 1.B). Finally, the modular states (MS) were obtained by applying community detection algorithm to the similarity matrix (Figure 1.C).

We propose two different frameworks i) “categorical” where the objective is to find the main modular structures over time, without any interest in their sequential order and ii) “consecutive” where the objective is to find the modular structures in a successive way.

### 3.1. Validation on simulated data

Figure 5 shows the results of the categorical method applied on the dynamic networks generated by the simulation scenario (*STD*_*noise*_ = 0.2). Four modular states were obtained: MS1, MS2, MS3 and MS4. Figure 5.A illustrates the modular states’ time courses, showing the most likely state at each time-window whilst Figure 5.B shows the 3D representation of the MS. Clearly, the four simulated modular structures have been successfully reconstructed. However, one time window that actually belongs to the background (i.e. random) has been wrongly affiliated to MS2. Moreover, MS3 state time course presented two false time window detection: one belongs to MS4, and the other belongs to the background. To quantitatively validate the obtained results, we compared the simulated structures (M1, M2, M3 and M4) to the reconstructed structures (MS1, MS2, MS3 and MS4) in terms of spatial and temporal similarities. The spatial similarities between the simulated and the reconstructed data are 0.99, 0.98, 0.99 and 0.98 for MS1, MS2, MS3, and MS4 respectively. The temporal similarities are 0.79, 0.83, 0.9 and 0.71 for MS1, MS2, MS3, and MS4 respectively.

**Figure 5.**
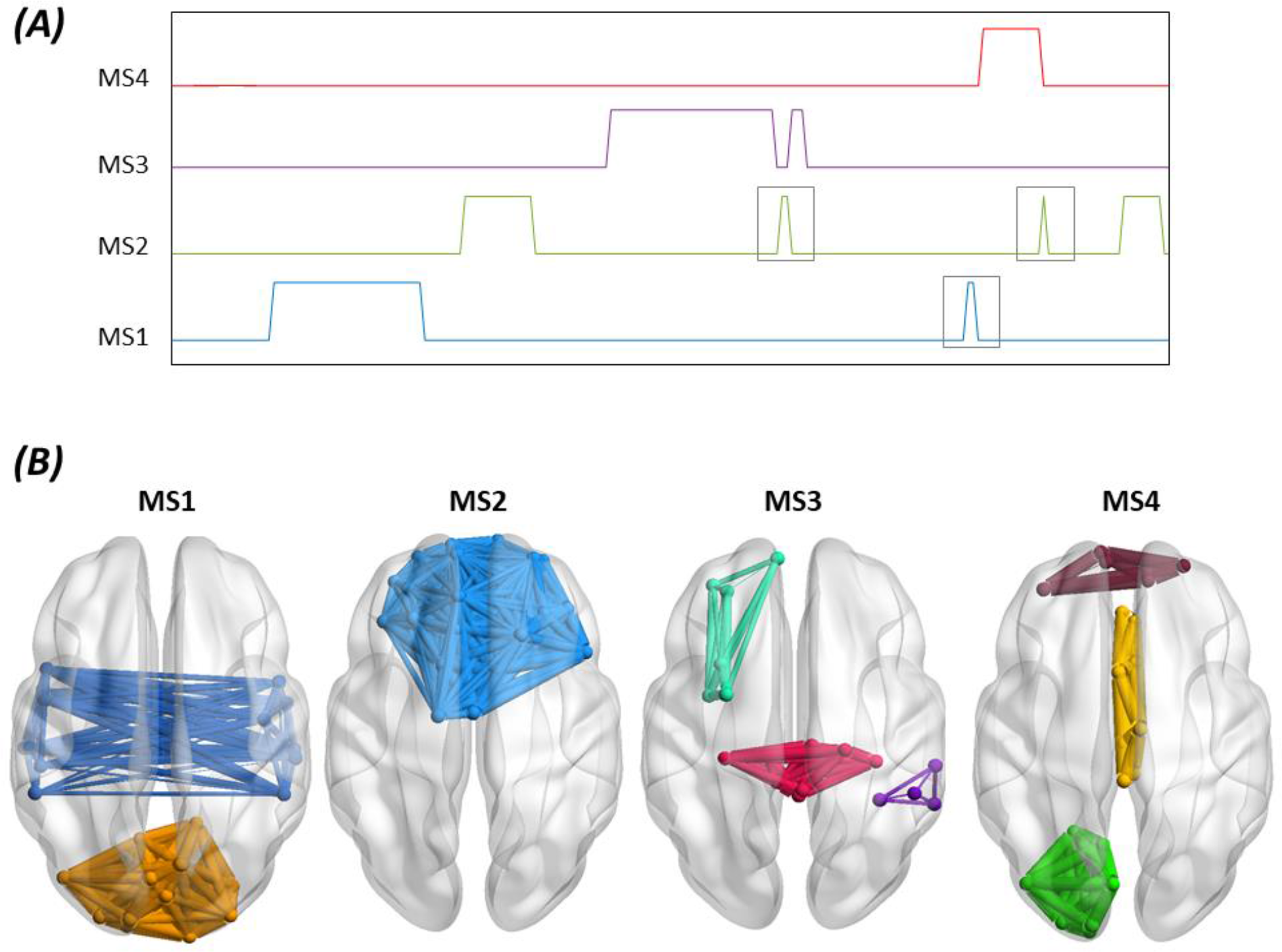
Results of the categorical method applied on simulated data. A) the time course of the four modular structures reconstructed. The grey square indicates false time-window detection. B) 3D representation of the four modular structures states.

Using the consecutive method, the algorithm has successfully segmented the similarity matrix yielding to the detection of five modular states (Figure 6.A). Their 3D representations are shown in Figure 6.B. One can remark that MS1 (spatial similarity=0.94; temporal similarity=0.88), MS2 (spatial similarity=0.99; temporal similarity=0.94), MS3 (spatial similarity=0.97; temporal similarity=0.77), MS4 (spatial similarity=1; temporal similarity=0.88), and MS5 (spatial similarity=0.95; temporal similarity=1) matched, temporally and spatially, the simulated networks generated at the corresponding time-windows.

**Figure 6.**
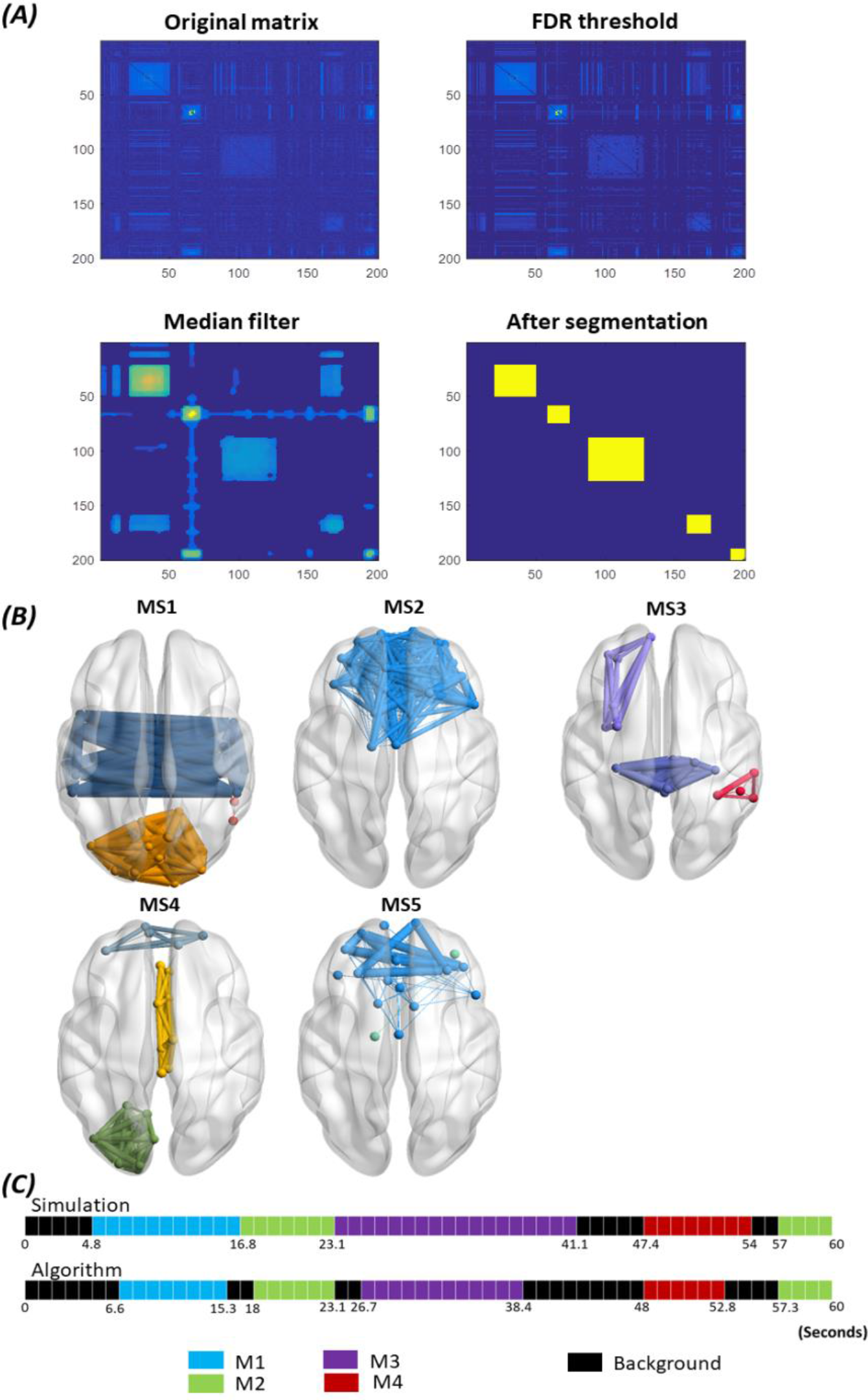
Results of the consecutive method applied on simulated data. A) The results of the segmentation algorithm used to derive the consecutive modular structures from the similarity matrix by: 1) Thresholding the matrix using FDR, 2) applying a median filter on the thresholded matrix and 3) extracting the most significant segments (See methods section for more details about the consecutive algorithm steps). B) The 3D representation of the five consecutive modular structures (MSs) obtained. C) The difference between the simulated time axis and the obtained time axis.

Results corresponding to *STD*_*noise*_ = 0.35; *STD*_*noise*_ = 0.5 are illustrated in the supplementary materials. In brief, results show that using the categorical algorithm, the spatial characterizations of the four modular states were successfully detected. However, the state time-course of MS3 failed to detect the second corresponding segment (Figure S1, S2). Using the consecutive algorithm, the five MS were temporally detected (Figure S3, S4).

### 3.2. Real data

#### 3.2.1. Dataset 1- *self paced motor task for healthy participants (MEG data)*

The consecutive algorithm was applied on the computed dynamic connectivity matrices averaged over all trials and subjects. The same dataset and methods was previously used in (O’Neill et al., 2017b, 2015; Vidaurre et al., 2016). The algorithm results in one significant MS found between - 0.5s and 1.5 s (Figure 7). As illustrated, this module implicates the sensory motor area, the post and pre-central regions of both hemispheres.

**Figure 7.**
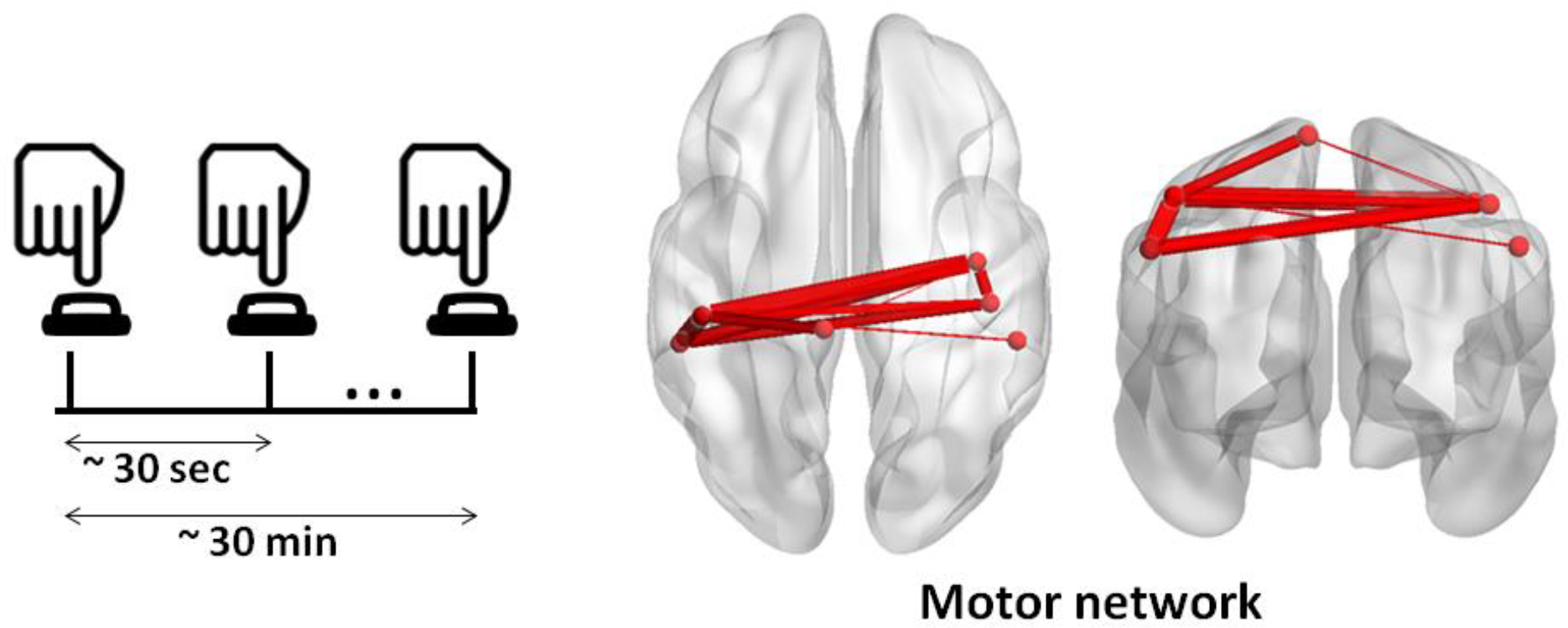
The MS of the MEG motor task obtained using the ‘consecutive’ method.

#### 3.2.2. Dataset 2- *picture naming task for healthy participants (dense-EEG data)*

The objective of using this dataset was to track the fast space/time dynamics of functional brain networks at sub-second time scale from the onset (presentation of the visual stimuli) to the reaction time (articulation). Hence, the consecutive version was applied on the dynamic connectivity matrices averaged over subjects.

Figure 8.A shows the obtained results revealing that the cognitive process can be divided into five modular structures: The first MS corresponds to the time period ranging from the stimulus onset to 130 ms and presents one module located mainly in the occipital region. The second MS is observed between 131 and 187 ms, and involves one module showing occipito-temporal connections. The third MS is identified between 188 and 360 ms, and illustrates a module located in the occipito-temporal region, and another module located in the fronto-central region. This structure was then followed by a fourth MS, found over the period 361-470 ms. MS4 was very similar to the previous MS but with additional fronto-central connections. The last MS is observed between 471 and 500 ms and it shows a module connecting the frontal, the central and the temporal regions. It is worth noting that these MS denote the transitions from the visual processing and recognition the semantic processing and categorization to the preparation of the articulation process (Hassan et al., 2015; VanRullen and Thorpe, 2001).

**Figure 8.**
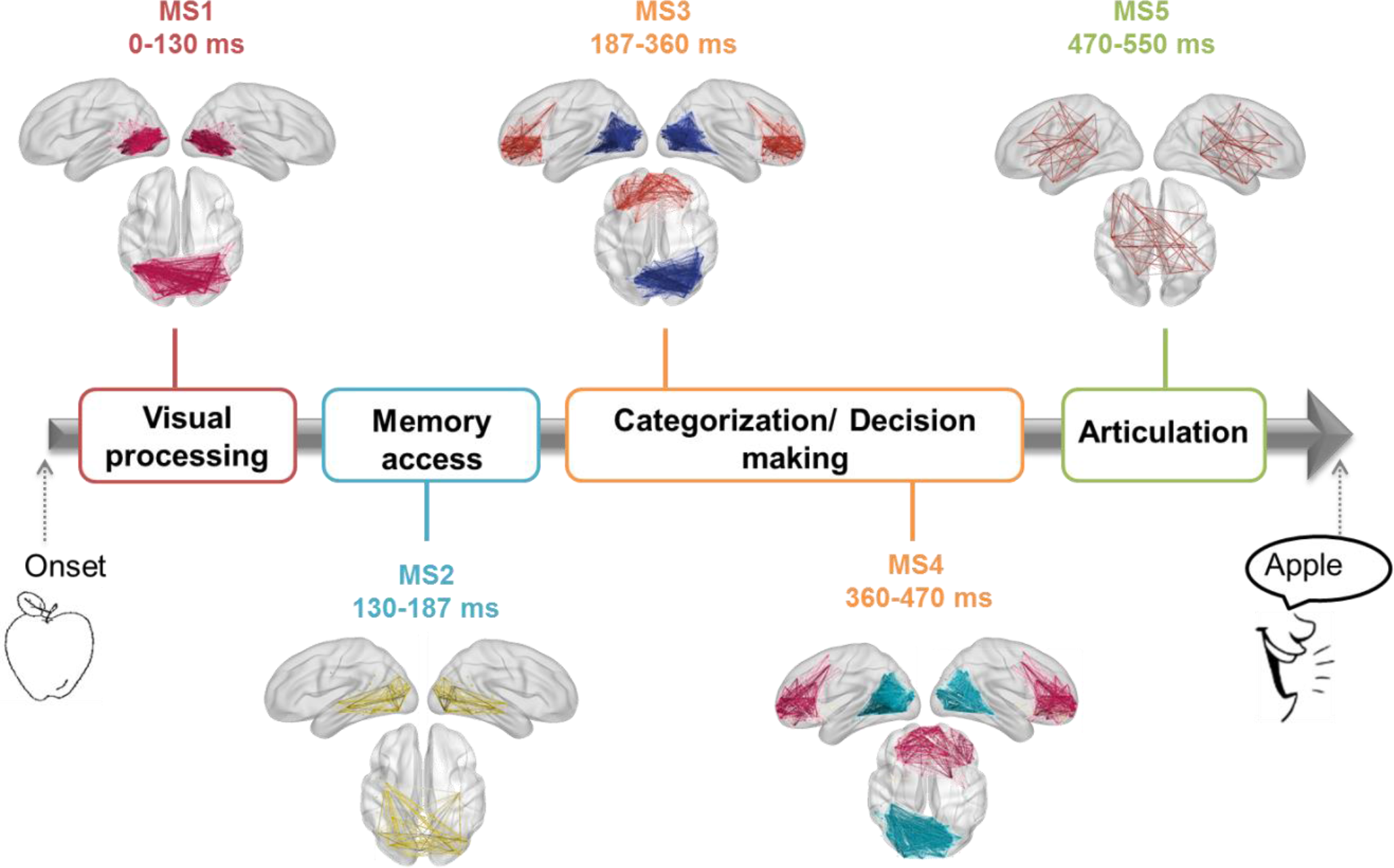
The sequential MSs of the EEG picture naming task obtained using the consecutive method and their corresponding cognitive functions.

#### 3.2.3. Dataset 3- *resting state in Parkinson′s disease patients (dense-EEG data)*

Our objective here was to validate the usefulness of the categorical version in detecting the modular alterations between G1, G2 and G3. To do that, the dynamic connectivity matrices of the three groups were concatenated over time, forming a single data tensor of dimension N x N x T, where N is the number of ROIs, and T is equal to the number of time-windows * the number of patients (Figure 9). The algorithm was then applied to identify the modular structures that are common to the three groups, and those who are specific to each group.

**Figure 9.**
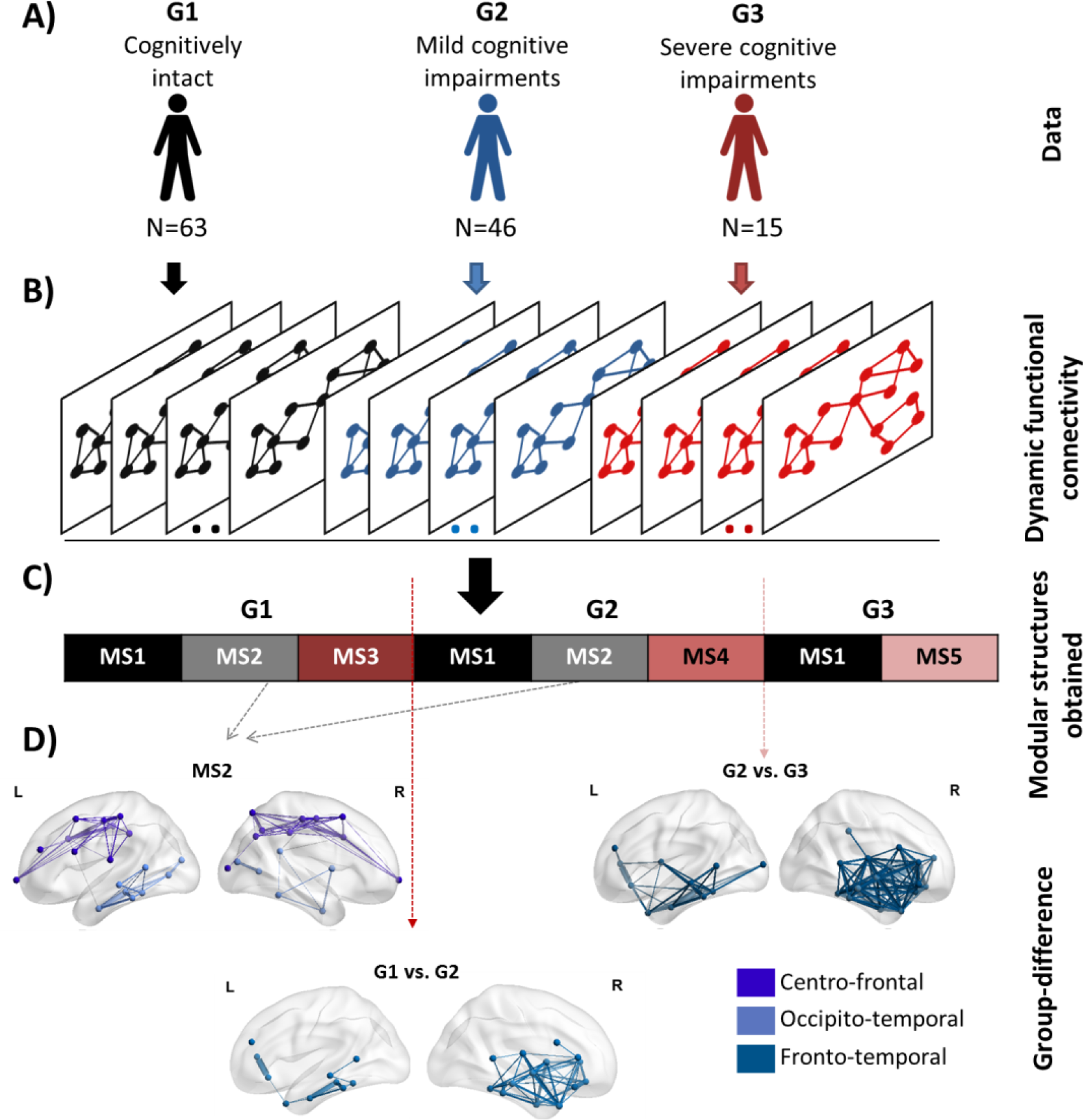
The analysis pipeline and the results of the categorical method applied on Parkinson’s disease EEG dataset. A) The dataset composed of 124 patients partitioned into three groups: G1) cognitively intact patients (N=63), G2) patients with mild cognitive deficits (N=46) and G3) patients with severe cognitive impairment (N=15). B) The functional dynamic connectivity matrices of the three groups concatenated over time. C) The five modular structures obtained after applying the categorical algorithm on the concatenated tensor. D) The modular differences between G1 and G2, G1 and G3, G2 and G3.

Results are illustrated in Figure 9. Five modular structures were identified (MS1, MS2, MS3, MS4 and MS5). Three MSs were found for G1 and G2. However, the number of MS decreased from three to two MSs in G3. Results revealed that MS1 was found to be present in the three groups while MS2 were present only in G1 and G2. The modular structure MS2 (absent in G3) is illustrated in Figure 9 and includes two modules involving mainly fronto-central and occipito-temporal connections. The difference between G1 and G2 was reflected by the absence of the structure MS3 replaced by the structure MS4 in G2. Results in Figure 9 showed that the difference concerns mainly the fronto-temporal connections. The difference between G2 and G3 was reflected by the absence of the structure MS4 from G2 and the presence of the structure MS5 in G3. Figure 9 showed that the functional disruptions between G2 and G3 are mainly fronto-temporal connections. It is worth noting that the fronto-temporal disruptions were widely reported in mild cognitive impairments (Beyer et al., 2007; De Haan et al., 2012; Song et al., 2011; H. Y. Zhang et al., 2009) while the central disruptions are widely observed in severe cognitive impairments (Hassan et al., 2017) and dementia (Song et al., 2011).

## 4. Discussion

In this paper, we have developed a novel framework to explore the fast reconfiguration of the functional brain networks during rest and task. This new method can be used to track the sequential evolution of brain modules during a task-directed paradigm or to identify the modular brain states that arise at rest. The simulation-based analysis showed the ability of the method to “re-estimate” the modular network structures over time.

The new framework was validated on three different EEG/MEG datasets i) MEG data recorded from 15 healthy subjects during a self-paced motor task, ii) Dense-EEG data recorded from 21 healthy subjects during a picture naming task and iii) Dense-EEG data recorded at rest from 124 Parkinson’s disease patients with different cognitive phenotypes. Results show that our method can track the fast modular states of the human brain network at sub-second time scale, and also highlights its potential clinical applications, such as the detection of the cognitive decline in Parkinson’s disease.

### 4.1. “Categorical” and “consecutive” processing schemes

The two processing schemes proposed here are both derived from the similarity matrix between the temporal modules (Step 1→3 in Methods section). However, each version highlights a specific characterization of the modular structures, which can be then exploited depending on the application (time/condition dependent). In particular, the results of the categorical algorithm on the simulated data reveal high spatial resolution and relatively low temporal resolution compared to those obtained using the consecutive algorithm. The low temporal resolution of the categorical version is reflected by the false (Figure 5) as well as the missed time-windows detection (Figure S1, Figure S2). In contrast, these time-windows were correctly detected by the consecutive version despite their short length (Figure 6, Figure S3 and Figure S4). Yet, the low spatial resolution of the consecutive version can be illustrated by MS5 (Figure 6, Figure S3, Figure S4) that should represent M2 (Figure 4). This is probably due to the categorical version using the maximum number of available data points to generate their corresponding MS, whilst the consecutive version treats each temporal segment solely.

We suggest using the consecutive version where sequential order of MSs is interesting to investigate such as the track of cognitive tasks. When the temporal aspects are not necessary, we would recommend the categorical version.

### 4.2. Tracking of fast cognitive functions

The brain dynamically reconfigures its functional network structure on sub-second temporal scales to guarantee efficient cognitive and behavioral functions (O’Neill et al., 2017a). Tracking the spatiotemporal dynamics of large scale networks over this short time duration is a challenging issue (Allen et al., 2014; Hutchison et al., 2013). In this paper, we aimed at examining how fast changes in the modular architecture shape the information processing and distribution in i) motor task and ii) picture naming task.

Concerning the self-paced motor task, it is a simple task where only motor areas are expected to be involved over time. Our results showed indeed that motor module is clearly elucidated related to the tactile movement of the button press. The spatial and the temporal features of the obtained module are very close to the significant component obtained by (O’Neill et al., 2017b) using the temporal ICA method.

The different MSs obtained in the EEG picture naming task are temporally and spatially analogous to the network brain states detected using other approaches such as K-means clustering by (Hassan et al., 2015). In particular, the first MS representing the visual network is probably modulated by the visual processing and recognition processes (Thorpe et al., 1996; VanRullen and Thorpe, 2001). The second MS reflects the memory access reflected by the presence of the occipital-temporal connections (Martin and Chao, 2001). In other words, the brain tries to retrieve the information related to the picture illustrated from the memory (Martin and Chao, 2001). In the third and the fourth MSs, we notice the implication of a separated frontal module. This module may be related to the object category recognition (tools vs. animals) and the decision making process (Andersen and Cui, 2009; Clark and Manes, 2004; Rushworth et al., 2011). After taking the decision, the speech articulation and the naming process is prepared and started (Dronkers, 1996). This is reflected by the MS5 that combines the frontal, the motor and the temporal brain areas.

### 4.3. Modular brain states and cognitive phenotypes in Parkinson’s disease

Emerging evidence show that Parkinson’s disease (PD) is associated with alteration in structural and functional brain networks (Fornito et al., 2015). Hence, from a clinical perspective, the demand is high for a network-based technique to identify the pathological networks and to detect early cognitive decline in PD. Here, we used a dataset with a large number (N=124) of PD patients categorized in three groups in terms of their cognitive performance: G1- cognitively intact patients, G2- patients with mild to moderate cognitive deficits and G3- patients with severe cognitive deficits in all cognitive domains. See (Dujardin et al., 2015; Hassan et al., 2017; Lopes et al., 2017) for more information about this database.

The obtained MSs presented in Figure 9 show that while some MSs remain unchangeable during cognitive decline from G1 to G3, others are altered and replaced by new MS. More specifically, the number of MSs detected in G3 has decreased compared to G1 and G2, MS3 in G1 was replaced by MS4 in G2 while MS4 in G2 was replaced by MS5 in G3. In addition, the alterations in G3 involve more distributed modules (central, fronto-temporal) than the alterations occurring between G1 and G2 (fronto-temporal modules) where the impairment still moderate.

Interestingly, the underlying modular differences between the MSs of groups are consistent with the previously reported studies that explored the networks changes in PD (Beyer et al., 2007; Bosboom et al., 2009; Haan and Pijnenburg, 2009; Hassan et al., 2017; Song et al., 2011; Y. Zhang et al., 2009). Particularly, the loss of fronto-temporal connections in PD is supported by several EEG and MEG studies (Bosboom et al., 2009; Hassan et al., 2017; Y. Zhang et al., 2009). Similarly, results of structural MRI studies reveal frontal and temporal atrophies in PD with mild cognitive impairment (Beyer et al., 2007; Song et al., 2011). Other functional (De Haan et al., 2012) and structural (Y. Zhang et al., 2009) studies showed that Alzheimer’s disease networks are characterized by fronto-temporal alterations. In addition, the brain regions involved in the modular alterations in G3 found in our study are in line with findings obtained by EEG edge-wise analysis (Hassan et al., 2017), and by structural MRI studies showing widespread atrophy associated with PD patients related dementia (Song et al., 2011).

### 4.4. Methodological considerations

First, we used a template anatomical image generated from MRIs of healthy controls for EEG/MEG source functional connectivity analysis. The template-based method is common practice in the absence of individual anatomical images and was previously employed by multiple EEG and MEG source reconstruction studies, because of non-availability of native MRIs (Hassan et al., 2017; Kabbara et al., 2018; Lopez et al., 2014). Furthermore, a recent study showed that there are few potential biases introduced during the use of a template MRI compared to individual MRI co-registration.

Second, the connectivity matrices in Dataset 3 (Parkinson’s disease analysis) were thresholded using a “proportional threshold” approach in contrast to other datasets (Picture naming and self-paced motor tasks) where a statistical threshold (FDR) approach was used (see Methods). The reason is that in dataset 3, three groups were compared. The proportional threshold approach, compared to other threshold approaches, ensures equal density between the analyzed groups (van den Heuvel et al., 2017). Moreover, studies suggest that FDR controlling procedures are effective for the analysis of neuroimaging data in the absence of inter-groups comparison (Bassett et al., 2013; Genovese et al., 2002; Patel and Bullmore, 2016).

Third, we are aware that spurious correlations caused by the problem of “source leakage” should be carefully considered. Here, we have adopted in each dataset the same pipeline (from data processing to networks construction) used by the previous studies dealing with the same dataset. Thus, for the MEG dataset, the correction for source leakage was performed by the symmetric orthogonalisation method proposed by (Colclough et al., 2015). Using the same pipeline also helps to avoid influencing factors caused by changing the source connectivity method, the number of ROIs, the connectivity measure or the sliding window length. By relying on previous studies (Hassan et al., 2017, 2015; O’Neill et al., 2016), we provide appropriate input -already tested and validated - to the algorithm, regardless of how they were obtained.

## 5. Acknowledgments

This work has received a French government support granted to the CominLabs excellence laboratory and managed by the National Research Agency in the “Investing for the Future” program under reference ANR-10-LABX-07-01. It was also financed by the Rennes University Hospital (COREC Project named conneXion, 2012-14). This work was financed by Azm center for research in biotechnology and its applications. GCO is funded by a Medical Research Council New Investigator Research Grant (MR/M006301/1). The study was also funded by the National Council for Scientific Research (CNRS) in Lebanon.

